# During haptic communication, the central nervous system compensates distinctly for delay and noise

**DOI:** 10.1101/2024.04.02.587670

**Authors:** Jonathan Eden, Ekaterina Ivanova, Etienne Burdet

## Abstract

Connected humans have been previously shown to exploit the exchange of haptic forces and tactile information to improve their performance in joint action tasks. As human interactions are increasingly mediated through robots and networks it is important to understand the impact that network features such as lag and noise may have on human behaviour. In this paper, we investigated the interaction with a human-like robot controller that provides similar haptic communication behaviour as human-human interaction and examined the influence and compensation mechanisms for delay and noise on haptic communication. The results of our experiments show that participants can distinguish between noise and delay, and make use of compensation mechanisms to preserve performance in both cases. However, while noise is compensated for by increasing co-contraction, delay compensation could not be explained by this strategy. Instead, computational modelling suggested that a feed-forward prediction mechanism is used to compensate for the temporal delay and yield an efficient haptic communication.

**Author summary:** Increasingly humans are making use of networks and robots to coordinate haptic interactions through teleoperation. However, with networks comes delays and noise that can change both the force that is transmitted and how we perceive that force. The haptic communication involved in joint actions, such as moving a piano or performing a pair spin, has been shown to improve performance, but how does delay affect this behaviour? We tested how participants tracked a moving target with their right hand when connected to a human-like robotic partner, when perturbed by delay or noise.

Through a comparison between noise and delay perturbation in experimental performance and in simulation with a computational model, we found that participants could from small values of perturbation identify if the perturbation was from delay or noise and that they adopted different compensation strategies in each case.

## Introduction

How do humans succeed in carrying out skilled motor tasks together, such as when moving a piano or when skaters perform a pair spin? The results of recent studies on joint tracking tasks [1–3] suggest that these collaborations are supported by the partners exchanging their motion plan via the haptic channel [4, 5]. Importantly, performance benefits arise between connected partners of different skills [1], where each partner must integrate information from their visual and haptic channels, and there is evidence that connected partners can have better task learning than when in a solo configuration [6, 7]. The ability to coordinate incoming sensory information from different modalities and with different time lags has been proposed to be critical for the central nervous system (CNS) to make sense of interactions with the environment [8, 9]. Here, the response to temporal delays is also of practical importance due to the transmission delays present when partners are connected by robotic systems, e.g. teleoperation in space applications with one partner on Earth and the other on a space station [10], or during remote physical training [11]. However, despite the physiological importance of sensorimotor delays and its noted affect on some sensory modalities [12]. how it influences haptic communication is not yet known. Therefore, we designed an experiment to investigate how haptic communication is affected by temporal delays present over digital connection [13].

This paper examines what mechanism physically connected individuals use to collaborate efficiently despite delayed haptic feedback. One possibility is that the CNS ignores the delay (*no compensation strategy*, Fig. 1A), in which case haptic communication (which can be modelled as in [4]) takes place as normal, such that performance should degrade with increasing delay. However, the CNS is known to inconspicuously fuse signals with different timing information [8], e.g., recognising the relationship between the visual signal of lightning and the delayed audio of thunder. This ability may extend to haptic communication, where sensor fusion occurs across partners [4]. Here, the evidence suggests that haptic communication arises due to the CNS identifying the interaction with the partner as task-relevant and optimally combining one’s own and the partner’s sensory information by considering their respective noise [4]. Corresponding to these results, the mismatch of delayed haptic information from the partner may instead be compensated as an additional noise (*compensation as noise strategy*, Fig. 1B). Alternatively, the CNS may be able to identify the delay and actively compensate for it, thus enabling the extraction of more specific information from the haptic signal (*compensation by delay prediction* strategy, Fig. 1C).

**Fig 1.**
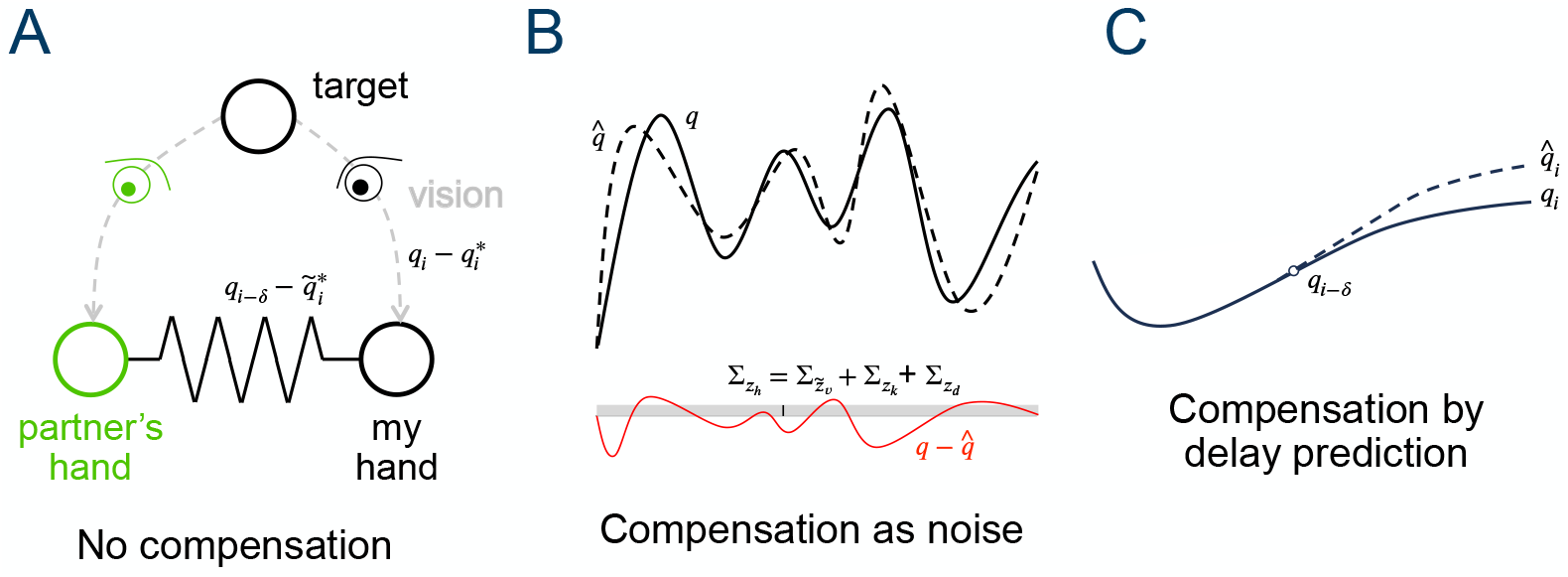
Possible neural mechanisms to deal with a delayed interaction with a partner. A: Haptic communication with *no compensation*, where the CNS understands that the haptic feedback is related to the visual task and uses it (without adjusting for delay) to infer the partner’s motion plan that is then combined with their own motion plan. B: *Compensation as noise* mechanism in which the CNS, additionally to A, considers the effect of the delay as an additional noise. C: *Compensation by delay prediction* mechanism in which the CNS, additionally to A, identifies the temporal delay and uses it to predict the current partner’s information.

To study how temporal delay affects haptic communication and to test these three possible strategies, we investigated how participants (connected to a human-like robot controller [5]) tracked a target moving along a multi-sine function with their dominant arm’s wrist flexion/extension. The participants moved an individual handle of the Hi5 dual robotic interface [14] to track a target with a cursor on their own monitor (Fig. 2A). They were connected by a virtual spring to the *robotic partner* of [4, 5], which induces similar interaction perception, behaviour and learning to a human [6, 15]. This enabled the systematic investigation of the effect of temporal delay and noise in haptic communication. We first reanalyzed the 20 participant’s data from [13] to investigate the effect of temporal delays up to 540 ms. To analyze whether the CNS compensates for delay as if it was noise (*compensate as noise*), we further tested the effect of noise in the haptic connection in a new group with another 20 participants. As muscle co-activation control is a known strategy to deal with noise [16, 17], the activity of a wrist flexor/extensor muscle pair was recorded. A questionnaire was used to compare perception of both types of perturbations. Experimental findings showed that while the noise data displayed a trend consistent with co-activation control, the delay data showed a different tendency. Therefore, we developed a computational model by extending the algorithm from [5] to investigate the mechanism used by the CNS to compensate for temporal delays and reproduce the observed experimental findings.

**Fig 2.**
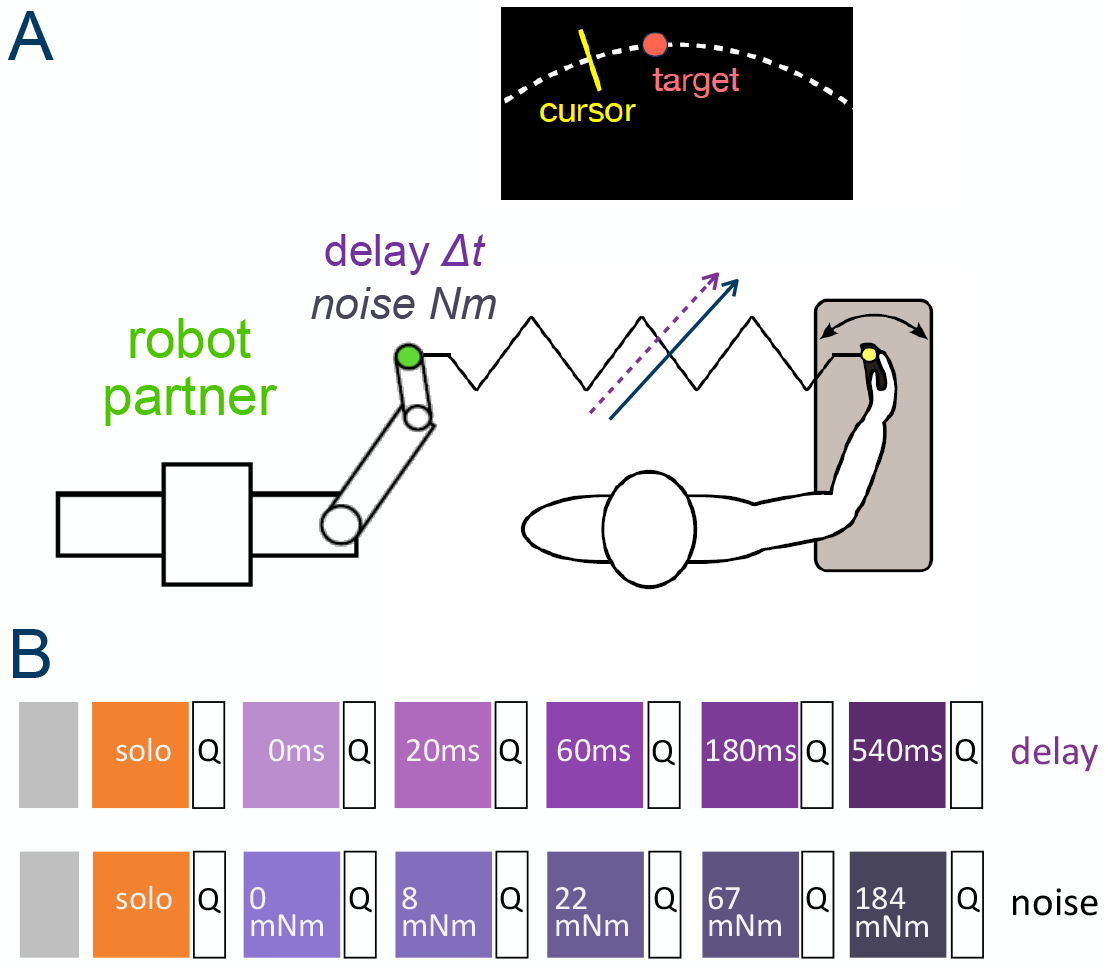
Experiment description. A: Participants tracked a randomly moving target with their wrist flexion/extension movement while being connected to a reactive robot partner (RP). B depicts the experimental protocol for the delay group [13] and noise group. The grey boxes represent the familiarisation/washout trials and the colourful blocks are the experimental conditions. Both groups started with the solo condition (i.e. without interacting with RP). They then were connected with a RP without delay/noise. Subsequently, the delay/noise was increased in every block. After each block, participants were asked to fill in a questionnaire. The order of the blocks was the same for all participants.

## Experimental results

The experiment consisted of six blocks of target tracking (Fig. 2B) conducted on the Hi5 robot with or without interaction with a *robot partner* (RP) (See Methods).

Participants were split evenly into two groups of 20, where each group was defined by the perturbation added to the interaction: *delay* ; *noise*. For each group, the initial block was a *solo* condition in which the participants tracked the target without any connection. This was followed by five additional blocks in which a connection to the robot was provided and a perturbation on that connection was increased after each block. Delays of {0,20,60,180,540} ms were added to the haptic feedback in the *delay group* and random noise torques with standard deviation {0, 8, 22, 67, 184} mNm were added in the *noise group*.

### Perception of delay and noise

After each experimental block, we asked participants to provide information about their perception of the haptic interaction by answering a questionnaire (see Supporting Information). In both the delay and noise groups, participants clearly identified the presence of forces in all interaction blocks (question: “During the task it seemed like I felt haptic forces”, Friedman test, delay: *χ*^2^(5) = 59.644, *p <* 0.0001; noise: *χ*^2^(5) = 57.847, *p <* 0.0001), such that the solo condition was distinguished from all other conditions (*p <* 0.05 for all pairwise comparisons to solo condition with post-hoc Wilcoxon tests).

The questionnaire also asked participants if they perceived delay and noise during each interaction block through a 5-point Likert scale from “strongly disagree” to “strongly agree” (see Fig. 3). In the delay group, participants appeared to clearly identify delay (Fig. 3A) even at small applied delay values, where the perception was always different to that of the no-delay condition (*χ*^2^(5) = 34.121, *p <* 0.0001; *p <* 0.05 for all pairwise comparisons between 0 ms and 20-540 ms delay groups). The delay group’s perception of noise (Fig. 3C) also changed depending on the applied delay (*χ*^2^(5) = 34.217, *p <* 0.0001). However, in contrast to their delay perception, the participants only had a higher perception of noise (relative to the no-delay condition) at the two highest applied delay levels, 180 and 540 ms (both *p <* 0.05).

**Fig 3.**
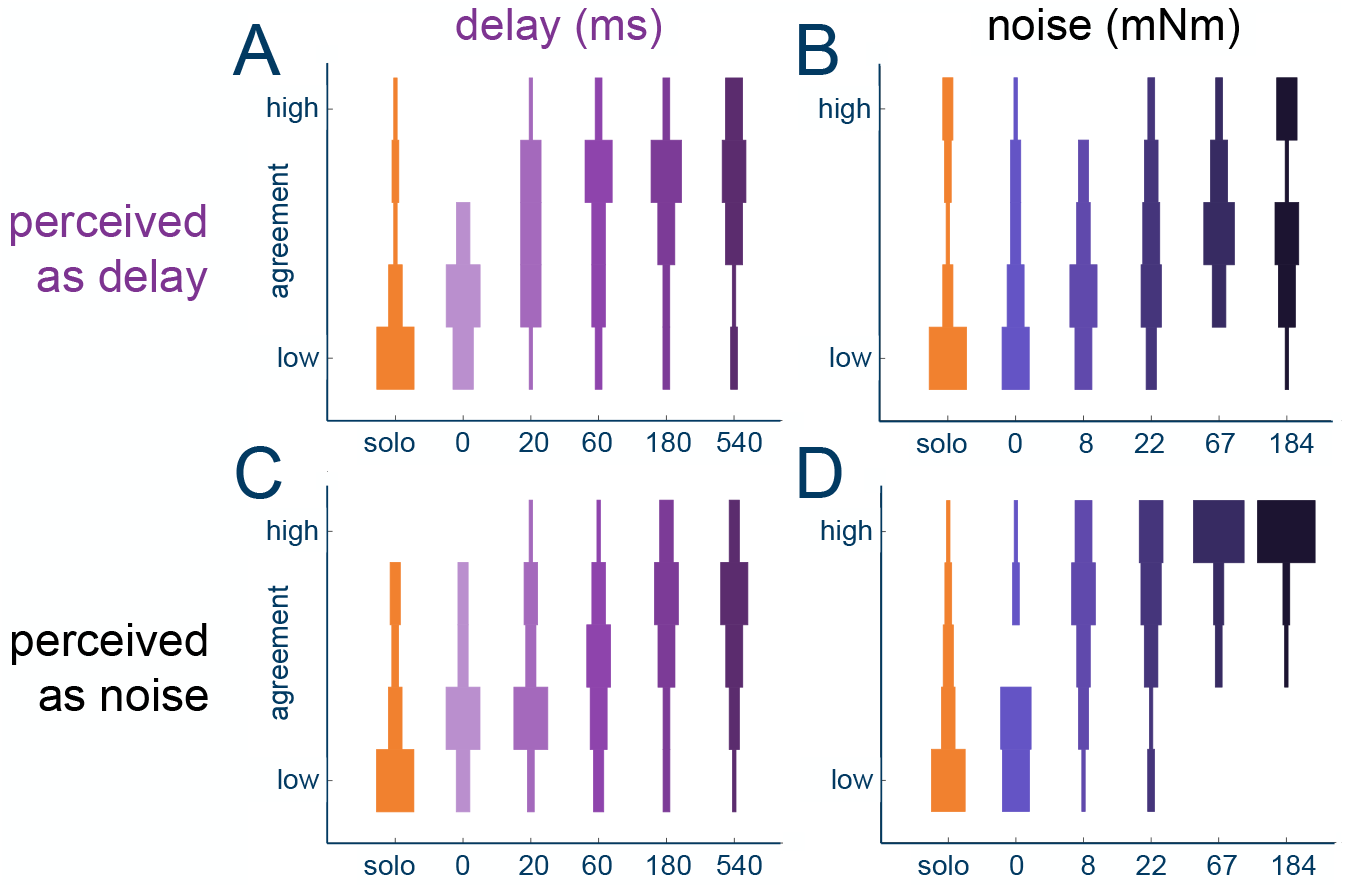
Perception during the interaction with a RP perturbed by delay or noise. (A) and (C) show the perception of delay and noise, respectively, for the delay group, while (B) and (D) show this perception for the noise group. Note the 5-point Likert scale goes from strongly disagree to strongly agree.

Different to the results in the delay group, participants in the noise group clearly identified the presence of noise from the smallest applied noise level (*χ*^2^(5) = 66.411, *p <* 0.0001; *p <* 0.01 for all pairwise comparisons between 0 mNm and 8-184 mNm noise conditions) as shown in Fig. 3D. In this group, the perception of the delay (Fig. 3B) also changed with the applied noise level (*χ*^2^(5) = 23.956, *p* = 0.00022), however, none of the conditions were found to be clearly different (all *p >* 0.05).

In summary, in both delay and noise groups, participants were able to recognise the presence of their respective perturbation from its smallest value. There was also some increase in perception of the non-adjusted factor in each group, however, there was limited confusion with a clear perception of the incorrect factor (compared to the solo condition) only reported in the delay group for its two highest delay levels. This indicates that the participants were able to perceive and distinguish delay from noise even without any knowledge of the experiment perturbation types.

### Different behaviours induced by delay and noise

We then analysed the participant’s performance (Fig. 4A) through their root mean squared tracking error. In both groups, the robot partner condition affected the performance (for delay: *χ*^2^(5) = 63.143, *p <* 0.0001; for noise:

**Fig 4.**
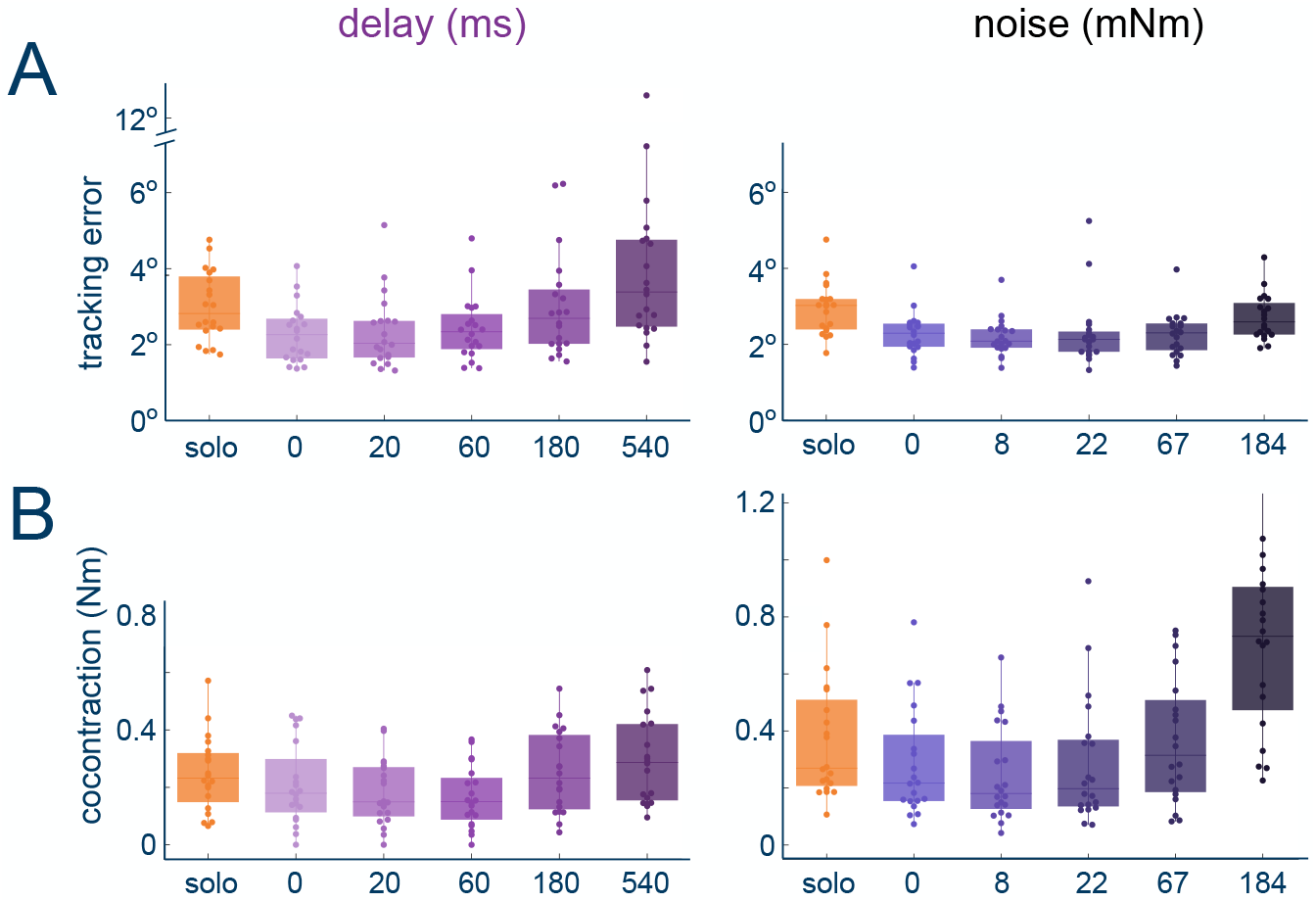
Performance and effort during the interaction with the robotic partner with delay (left) or noise (right) perturbation: tracking accuracy (A) and co-contraction (B). Each dot represents the average value in each block for one participant.

*χ*^2^(5) = 62.257, *p <* 0.0001). Here, as has been previously observed [4, 5], in both groups the addition of the robot partner (and the potential for haptic communication) improved the participant performance (solo vs. 0 for delay:

*W* = 210, *Z* = 3.9199, *p <* 0.0001; and noise: *W* = 210, *Z* = 3.9199, *p <* 0.0001).

However, while the participants were always able to compensate for the perturbation in the noise group such that the performance was always better to that of the solo condition (all *p <* 0.04) and not distinguishable from the condition working with a RP without perturbation (*p >* 0.05 for pairwise comparisons between 0-noise and 8,22,67 mNm), this was not the case for the delay group. Instead, here although the participants’ performance was not clearly affected by small applied delays (0 vs. 20 ms: *W* = 82, *Z* = −0.85865, *p <* 0.4091; 0 vs. 60 ms: *W* = 61, *Z* = −1.6426, *p* = 0.3162), they were not able to compensate for the larger applied delays (0 vs. 180 ms: *W* = 23, *Z* = −3.0613, *p* = 0.0073; 0 vs. 540 ms: *W* = 3, *Z* = −3.8079, *p <* 0.0001) such that at 540 ms their performance was worse than that of the solo condition (solo vs. 540 ms: *W* = 31, *Z* = −2.7626, *p* = 0.0211).

To understand if participants were using the previously observed co-activation compensation strategy [18] to aid in their response to the applied perturbations, we analysed their co-contraction (Fig. 4B) computed as the minimum measured absolute muscle activity of the flexor and extensor muscles over a trial (see Methods).

Participants changed their co-contraction in response to both applied perturbation types (delay: *χ*^2^(5) = 40.829, *p <* 0.0001; noise: *χ*^2^(5) = 58.6, *p <* 0.0001). However, while the noise group displayed a trend of increasing the median co-contraction with each subsequent applied perturbation level after the 8 mNm condition such that the co-contraction at the highest applied noise level was clearly larger than the no delay co-contraction (0-184 mNm: *W* = 0, *Z* = −3.9199, *p <* 0.0001), this was different in the delay group. Instead here, the participants held their co-contraction levels roughly constant for all small applied delay values (*p >* 0.05 for all pairwise comparisons between solo, 0 ms, 20 ms and 60 ms conditions except for solo-60 ms:

*W* = 25, *Z* = −2.9866, *p* = 0.0169, where the co-contraction was slightly lower). There was then an increase for the two highest applied delay values (60 ms-180 ms:

*W* = 1, *Z* = −3.8826, *p <* 0.0001; 60 ms-540 ms: *W* = 0, *Z* = −3.9199, *p <* 0.0001), in which the co-contraction level was not clearly different between the two levels (*W* = 25, *Z* = −2.9866, *p* = 0.0169). In contrast to the noise group, this increase in co-contraction did not prevent an increase of tracking error.

Finally, we examined if the two different perturbations affected the user trajectories, through the cross-correlation delay between the reference and user trajectory (Fig. 5A), and the motion smoothness (Fig. 5B) as computed by the SPARC measure [19]. The addition of the robot partner and perturbations affected the cross-correlation delay of both groups (delay: *χ*^2^(5) = 36.564, *p <* 0.0001; noise: *χ*^2^(5) = 48.724, *p <* 0.0001) where interestingly working with the robot partner without perturbation resulted in a reduced cross-correlation delay (both *p <* 0.0001). For the delay group, the cross-correlation delay in high-delay conditions was higher compared to smaller delay conditions (20 - 540 ms: *W* = 20, *Z* = −3.1733, *p* = 0.0007; 60 - 540 ms: *W* = 11, *Z* = −3.5096, *p* = 0.0005; 60 - 180 ms: *W* = 5.5, *Z* = −3.6028, *p* = 0.0003) suggesting that at those delay conditions participants struggled to counteract the delay. In contrast, for the noise group, the cross-correlation delay was higher in the solo and 0-noise condition (*p <* 0.0001 for all pairwise comparisons with solo; 0-8 mNm:

**Fig 5.**
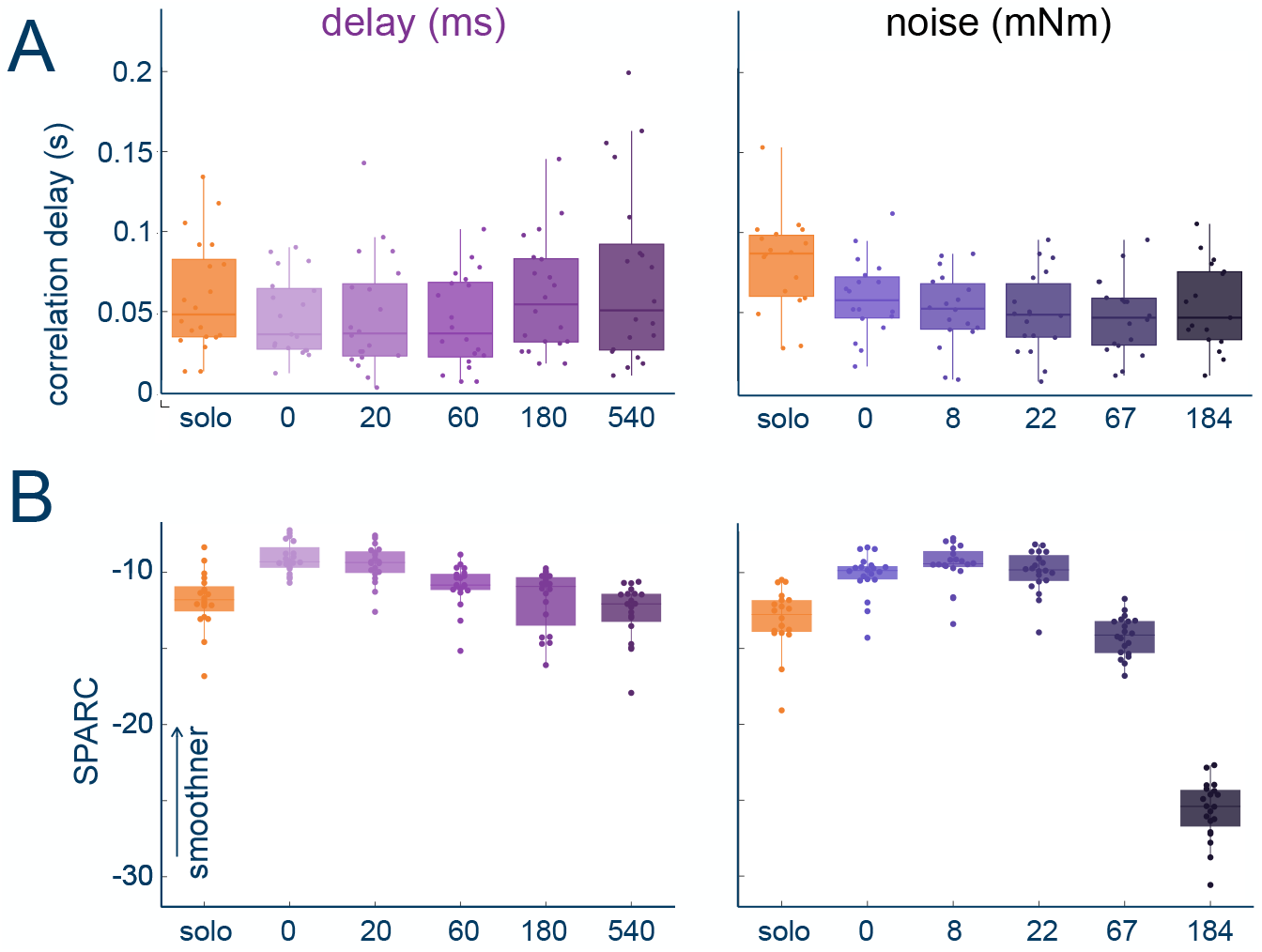
Lag and noisiness during the interaction with the robotic partner with a delay: cross-correlation delay (A) and smoothness (B). Each dot represents the average value in each block for one participant.

*W* = 147, *Z* = 2.6787, *p* = 0.0437; 0-22 mNm: *W* = 148, *Z* = 2.7222, *p* = 0.0366; 0-67 mNm: *W* = 163, *Z* = 3.3752, *p* = 0.0019) and all other conditions were similar to each other (*p >* 0.05 for all other pairwise comparisons).

The participants smoothness was also affected by the perturbation (delay: *χ*^2^(5) = 75.771, *p <* 0.0001; noise: *χ*^2^(5) = 93.486, *p <* 0.0001), where as has been previously observed [15] the addition of the robot partner without perturbation resulted in smoother motion (both *p <* 0.0001). In both cases the high perturbation levels were different to that of the no perturbation robot partner (delay: *p <* 0.05 for 0-180 ms and 0-540 ms; noise: *p <* 0.05 for 0-67 mNm and 0-184 mNm).

These results confirm that haptic communication improves interaction performance. While this performance is then affected by both the delay and noise perturbations, the effect is distinctly different in the two groups. For the noise group, participants appear to compensate as would be predicted by the co-activation compensation strategy, where their co-contraction increases with the applied perturbation level, and while there is an observed degradation of motion smoothness, there is no tracking performance that is worse than that of the solo condition. In contrast for the delay group, while there is an increase in co-contraction for the two highest applied perturbation levels, this increase does not result in compensation for the performance. Instead, participants appear to compensate for the applied delay where only at the highest level of injected delay was there a different lag to the other delay perturbation levels, and this was still not different from their own lag in the solo condition.

### Simulation results

The experimental results suggest that the response of the noise group is consistent with that predicted by the co-activation compensation strategy. However, the delay group behaviour was not consistent with these predictions. Could the results be explained strictly from the interaction mechanics, without any compensation? To evaluate this, we simulated the experimental scenario (with the same number of trials and blocks) of a participant being connected (by a virtual spring) to a partner with delayed haptic feedback. Here both partners were simulated using the model of haptic communication presented in [4, 5], which is based on four principles: i) The CNS of each participant is able to identify that the haptic feedback that they receive is related to the (visual) tracking task; ii) By using a model of the partner’s control, the CNS is able to extract their motion plan; iii) The participants then combine their own and partner motion information in a stochastic optimal way, yielding Bayesian sensor fusion of the visual and haptic information; iv) The elasticity of the haptic connection to the partner is considered as an additional source of noise. To simulate the effect of the delayed haptic feedback, we delayed the virtual spring torque for one of the participants with a delay value matching those of the experimental condition.

When both partners were modelled as in [5] such that there was *no compensation*, the performance (Fig. 6C) did not match that observed in the experimental data (Fig. 6A). Here, while there was little difference for small applied delay values, the performance could became unstable at larger delay levels such that it overshot the participant performance. Therefore we conclude that the participants had a mechanism to compensate for the delay.

**Fig 6.**
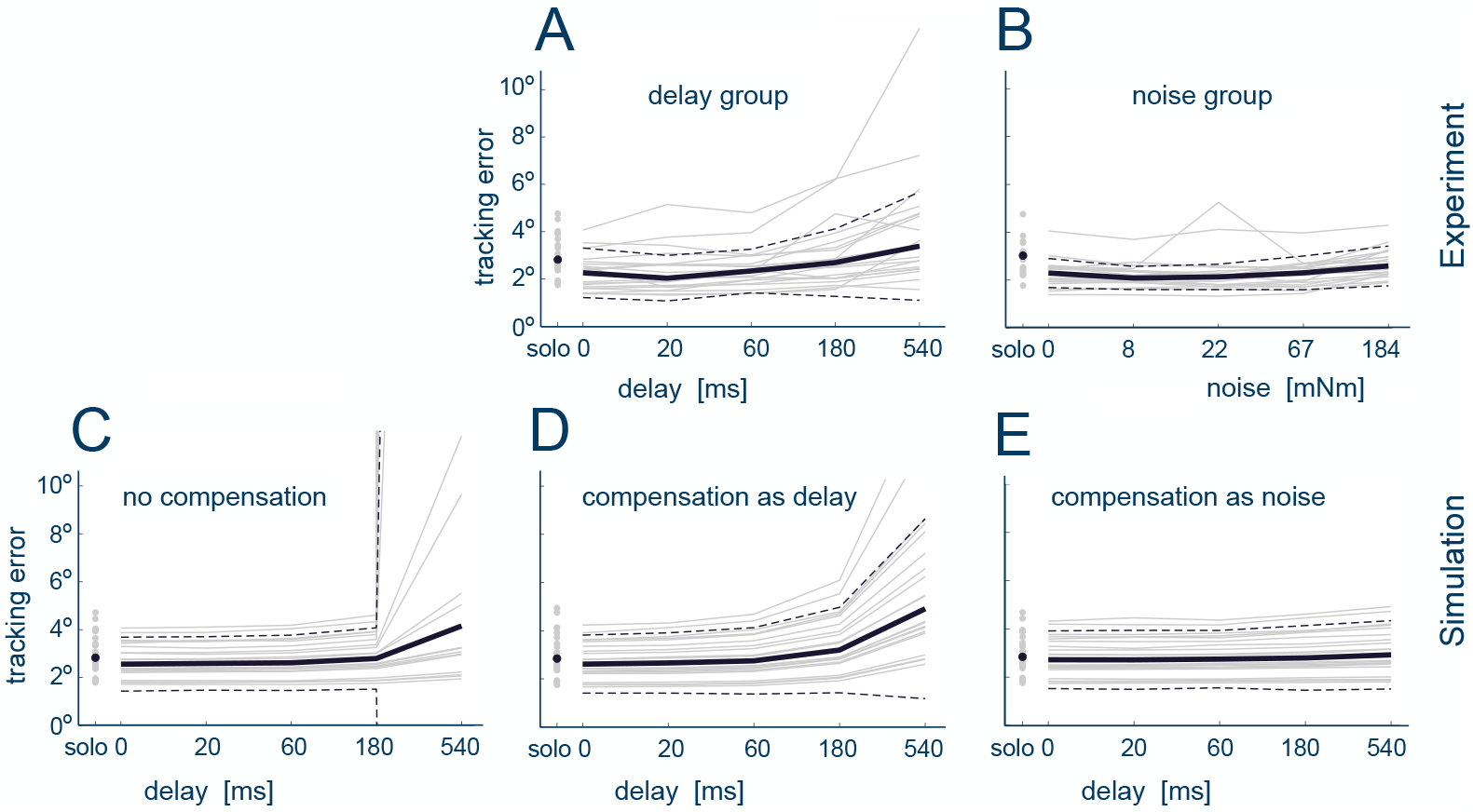
Tracking performance in the experiment and simulation to analyze the mechanism to compensate for temporal delays. Experimental results for the delay (A) and noise (B) groups. Simulated results for delay compensation set as: no compensation (C), compensation as noise (D) and compensation by delay prediction (E).

Participants did appear able to recognise delay (Fig. 3A). However, they may have adopted different strategies for its compensation which could lead to the observed motion characteristics. To investigate this compensation mechanism we extended the model developed in [5] to consider *compensation by delay prediction* as well as the *compensation as noise* (see Methods). In the *compensation as noise* simulation, this consisted of modifying iv) such that both the elasticity of the haptic connection and the delay of the feedback of that connection were considered as additional sources of noise (Fig. 1B). In the *compensation by delay prediction* simulation, it was instead assumed that as a part of i) and ii) the participant would be able to identify the delay and use their model of the interaction and delay to predict the future value for the haptic feedback (Fig. 1C).

The simulation results exhibit clear differences between the two models where similar to what was observed for the noise group experimental results, the *compensation as noise* incorrectly predicted the delay response performance (Fig. 6D) to be roughly constant across the different delay perturbation levels. In contrast, the *compensation as delay prediction* was able to produce a similar trend of results matching the similar performance characteristics with small delay as well as the error growth as the delay value increased (Fig. 6E).

## Discussion

This paper investigated the mechanism of haptic communication during a tracking task in response to delay and noise perturbations. Our results indicate that participants can still exploit haptic communication in the presence of both small delays and moderate levels of noise, where haptic communication resulted in improved smoothness, accuracy and correlation delays. Interestingly, the participants were able to correctly recognise the presence of delays and noise from their smallest values with limited confusion between the correct perturbation and other possible perturbing modalities. They then appear to compensate for each of these two different perturbation types with different strategies.

In response to increased noise, our findings indicate that while participants had a clear reduction of smoothness, their co-contraction showed a trend to increase with increasing noise, as would be predicted by the co-activation compensation strategy [16, 18]. The delay group instead showed limited increase in co-contraction. Here, the results appear instead to be consistent with a strategy of *compensation by delay prediction*, where the CNS would identify the delay and compensate for it when predicting the target movement. Such a strategy is consistent with recently observed findings for the multisensory integration of information across the visual and haptic channels [20] and suggests that such an integration extends to the scenario of haptic communication in which the user is integrating feedback from a partner. It is worth noting the two applied delay levels for which noise was perceived (delays of 180 and 540 ms) both also coincided with the predicted increase in co-contraction. It is therefore possible that while participants used *compensation by delay prediction*, in response to their insufficient compensation they identify noise and then try to compensate (ineffectively) through co-activation.

While our findings showed similar tracking error and smoothness improvements resulting from haptic communication as had been previously observed [1, 4], we further observed a reduction in the cross-correlation delay that had not been previously considered. This supports the understanding that the participant is able to incorporate the information within the haptic channel in order to improve their own motion plan. The reduced lag then suggests that this information not only improves the quality of their estimation but also aids them to adapt quicker then what they normally would.

In summary, physical connections over the haptic channel can improve user performance and is robust to small to medium sized perturbations in the form of delays and noise. This compensation is made possible by the participant being able to uniquely perceive the presence of delay or noise and then compensate specifically to these different perturbations.

## Methods

### Participants

The experiment was approved by the Research Ethics Committee of Imperial College London and carried out by 40 participants (21 female, 19 male) without known sensorimotor impairment aged 24.02 *±* 3.19 years old. Each participant gave informed consent, and filled in a demographic questionnaire as well as the Edinburgh handedness form [21] before starting the experiment. All participants performed the task with their dominant hand, where four participants were left-handed. Participants were divided into two groups that each experienced only one type of perturbation: human-robot interaction with i) time delay (data collected in [13]) or ii) with haptic (torque) noise.

### Experimental setup and protocol

The experimental task was designed to replicate the tasks of existing studies of haptic communication [15]. Two participants completed the experiment at the same time, with their dominant arm attached to the Hi5 dual robotic interface [14]. The participants were instructed that within the experiment that they might interact with a robot, another human, or on their own. They were then visually separated from one another by a curtain and each participant’s wrist flexion/extension movement interacted with the Hi5 interface, which was controlled at 1000 Hz, while the wrist angle data was recorded at 100 Hz.

Each participant was asked to track a moving target “as accurately as possible” using the wrist flexion/extension of their dominant hand (Fig.2A). The target trajectory was given by

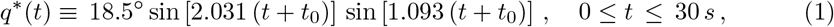

where to minimize the learning of the trajectory each trial started from a randomly selected starting time {*t*_0_ ∈ [0, 30] s | *q*^∗^(*t*_0_) ≡ 0}. The participant’s wrist flexion/extension was connected to a *robotic partner* (RP) with angle *q*_*r*_ through a virtual elastic band described (in Nm) by

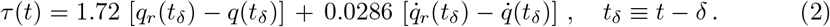

The robotic partner (RP) is a reactive controller [4, 15] that mimics human interaction behaviour, including differences in skill level. This is achieved through a sensory augmentation approach in which the information coming from the haptic connection is used to infer the partner’s target [4], which is then combined with their own target in a stochastically optimal manner.

For eq. (2) the damping and stiffness constants were chosen to match the conditions of medium stiffness in [4], with which an interaction with a interactive agent was clearly perceived by participants [15]. In the *delay group δ* ∈ {0, 20, 60, 180, 540} ms was used for the delayed interaction torque (while the robot partner received the torque without delay). For the *noise group δ* = 0 while the torque was perturbed by Gaussian noise *ν*:

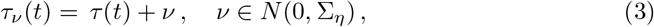

where the standard deviation Σ_*η*_ ∈ {7.5,22.5,66.7,184} mNm was used.

Surface electrodes were used to record electromyographical (EMG) activity from the wrist flexor carpi radialis (FCR) and extensor carpi radialis longus (ECRL) muscles. This was calibrated through a process in which participants were asked to flex/extend while their wrist was locked by the device at 0^*°*^ corresponding to the participant’s most comfortable position. The participant was asked to produce flexion and extension torques of {1, 2, 3, 4} Nm for 2 seconds, first flexion then extension, with a rest period of 5 seconds between each activation to prevent fatigue. This EMG data was linearly regressed with the measured torque to estimate the relationship between muscular activity and torque. Then the *co-contraction* was computed as

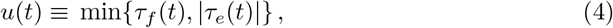

where *τ*_*f*_ (*t*) ≥ 0 and *τ*_*e*_(*t*) ≤ 0 are the flexor and extensor torques, computed from the respective EMG signals. The average co-contraction over all participants was computed from each participant’s normalised co-contraction, calculated as

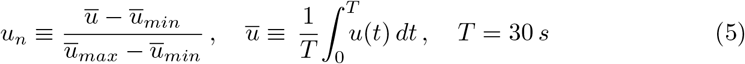

with ū _*min*_ and ū _*max*_ the minimum and maximum of the means of all trials of the specific participant.

The experiment protocol is described in Fig.2B. In the initial solo block, each participant attempted five trials of the task without a haptic connection to be familiarised with the task and to minimise subsequent learning effects. In the main experiment, participants carried out six blocks, each of ten trials. Each block included seven experimental trials followed by three washout trials of the solo condition. The first experimental condition was without any interaction and in the following five blocks assistance from a robot partner was introduced for both delay and noise experiments. The RP’s motor noise deviation was set after the solo block as equal to the deviation observed in the participant’s tracking movement during the final solo trial, to ensure that the participant and the RP have similar skill level [4]. This was chosen as it has been found to provide the best performance [1]. Each trial took 30 s and was followed by a 5 s break.

The delay and noise within the robotic assistance trials were increased from each block to the next with values {0, 20, 60, 180, 540} ms for delay and {0, 8, 22, 67, 184} mNm for noise. The sequence of the blocks with increasing level of perturbation was identical for each participant within delay and noise experiments. The delay levels were chosen to include small delay values considered in [22] and values greater than the threshold for performance loss found in [23]. The Gaussian noise values were then set to approximate the effect of the delay. Here the noise torque standard deviation Σ_*η*_ was set so that three standard deviations was equivalent to the likely maximum error torque caused by a given delay, which was approximated by 1.72 *·* max{*q*^∗^(*t*) − *q*^∗^(*t* − *δ*)}. After each block, the participants had to answer questions about their perception of the interaction (see Supporting Information).

### Simulation framework

We evaluated the mechanism for delay using the computational model developed in [5]. In this discrete time model, the control of the wrist is modelled as a linear controller that acts on a double integrator system which describes the wrist angle *q*. The state space dynamics at time-step *i* are given by

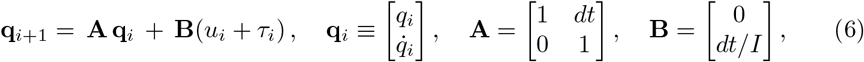

where *dt* is the time differential and *I* = 0.002 kg m^2^ the wrist’s moment of inertia (as defined for experiments with the Hi5 robot in [5]). The control input *u*_*i*_ is determined by the linear feedback control law

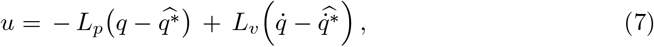

where 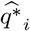 denotes the participant’s estimate of the trajectory. *L*_*p*_ and *L*_*v*_ are the proportional and derivative gains determined to minimise a quadratic cost function of error and effort [16]. Moreover, the system is influenced by the haptic interaction torque *τ*_*i*_. This torque is set to 0 during solo trials. It instead acts as a spring and damper torque and is set as in eq. (2) for interaction trials connecting the agent to their partner whose dynamics similarly evolves with a form given by eq. (6).

The key feature of the model is that the partners improve their estimate 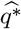 through visual feedback and haptic information from the interaction. Here, their own target information is combined with the partner’s target information 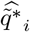, as determined from the interaction force. This integration of the partner’s target information is carried out through a Kalman filter in which the measurement **z**_*i*_ is given by

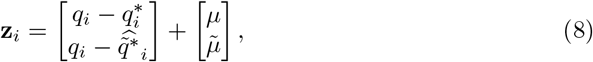

where 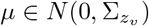 and 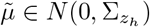. This assumes that 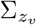 characterises the visual noise naturally present in the participant’s tracking, while 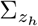 characterises the haptic noise and is composed of the partner’s visual noise 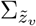 and additional noise resulting from the virtual band elasticity 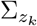 such that 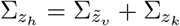.

### Delay compensation strategies

Three delay compensation strategies are considered within the simulation: i) no compensation; ii) compensation as noise; iii) compensation as delay prediction. In the *no compensation* strategy participants were simulated to directly use above interaction model with a delayed estimation of the partner’s error, i.e.

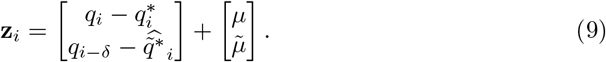

In the *compensation as noise* strategy participants were modelled to use the same delayed measurement as in eq. (9). However, they were assumed to consider the haptic signal to be noisier with an additional noise 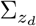 associated to the given delay level such that

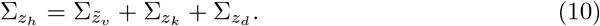

The magnitude of this additional delay generated noise was determined by simulating the participant’s solo performance with the delay and then mapping the resulting error to a noise value through the error to noise regression determined in [5].

Finally, in the *compensation as delay prediction* strategy the participants were instead modelled to be able to identify both the presence of the delay as well as its magnitude. They were then assumed subsequently adjust their haptic measurement through forward integration with the known system dynamics eq. (6) and visual target sequence. Here the current error estimate 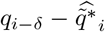 was updated by iteratively applying eq. (6) (without the unmeasured torque *τ*_*i*_) for the number of delayed time steps (given as *δ*/*dt*).

### Data Analysis

To investigate the delayed force exchange’s effect on participant performance, the Root-Mean-Square Error (RMSE), the smoothness metrics SPARC [19], the cross-correlation delay and the co-contraction were analysed. The *cross-correlation* delay corresponds to the time interval between the target’s movement and the participant’s resulting motion and was calculated as the time lag at which the cross-correlation between the target and participant’s positions was the highest. To understand how participants perceived the changes in delay, a questionnaire composed of a 5-point Likert-scale items was analysed (see the question list in Supporting Information).

Since we could not compare the delay and noise magnitudes directly, a separate analysis was conducted for each perturbation group and the group tendencies were then qualitatively compared. Since each metric was found to not be normally distributed, the influence of the perturbation on each metric was explored through Friedman tests.

Post-hoc analysis between individual perturbation levels was conducted using a paired Wilcoxon sign-rank test with the Hommel adjustment to control the family-wise error rate. For each objective values (RMSE, SPARC, cross-correlation delay, co-contraction) the analysis was conducted for each participant using the averaged value over all trials in a block.

## Supporting Information

The full list of the questions of the questionnaire used in the study after each experimental block, where questions (1-12) had response options *{*1/5: strongly dis/agree*}* [13]:

1. I was controlling the movement I saw.
2. I was the one who produced the movement I saw.
3. It seemed like I felt haptic forces.
4. It seemed like I felt haptic perturbation.
5. It seemed like I felt haptic interaction with an agent.
6. It seemed like I felt haptic noise.
7. It seemed like I felt haptic feedback.
8. It seemed like I felt haptic assistance.
9. It seemed like I felt haptic resistance.
10. It seemed like I felt haptic interaction with a delay.
11. It seemed like I felt that my hand was heavier.
12. It seemed like I felt no haptic feedback of any kind.
13. Was the interaction predictable? {1/5: un/predictable}
14. Was the interaction natural? {1/5: natural/artificial}
15. Was the interaction disturbing or helpful? {1/5 disturbing/helpful}
16. Was the interaction useful? {1/5 useful/harmful}

## Acknowledgements

This work was supported in part by the EU H2020 grant ICT-871803 CONBOTS and by the UK EPSRC EP/R026092/1 FAIRSPACE program.

## References

1. Ganesh G, Takagi A, Osu R, Yoshioka T, Kawato M, Burdet E. Two is better than one: Physical interactions improve motor performance in humans. Scientific Reports. 2014;4:3824.

2. Beckers N, van Asseldonk EH, van der Kooij H. Haptic human–human interaction does not improve individual visuomotor adaptation. Scientific Reports. 2020;10(1):1–11.

3. Mojtahedi K, Whitsell B, Artemiadis P, Santello M. Communication and inference of intended movement direction during human–human physical interaction. Frontiers in Neurorobotics. 2017;11:21.

4. Takagi A, Ganesh G, Yoshioka T, Kawato M, Burdet E. Physically interacting individuals estimate the partner’s goal to enhance their movements. Nature Human Behaviour. 2017;1(3):0054.

5. Takagi A, Usai F, Ganesh G, Sanguineti V, Burdet E. Haptic communication between humans is tuned by the hard or soft mechanics of interaction. PLoS Computational Biology. 2018;14(3):e1005971.

6. Ivanova E, Eden J, Carboni G, Krüger J, Burdet E. Interaction with a reactive partner improves learning in contrast to passive guidance. Scientific Reports. 2022;12(1):15821.

7. Batson JP, Kato Y, Shuster K, Patton JL, Reed KB, Tsuji T, et al. Haptic coupling in dyads improves motor learning in a simple force field. In: International Conference of the IEEE Engineering in Medicine & Biology Society (EMBC); 2020. p. 4795–4798.

8. Parise CV, Ernst MO. Correlation detection as a general mechanism for multisensory integration. Nature Communications. 2016;7(1):1–9.

9. Cao Y, Summerfield C, Park H, Giordano BL, Kayser C. Causal inference in the multisensory brain. Neuron. 2019;102(5):1076–1087.

10. Imaida T, Yokokohji Y, Doi T, Oda M, Yoshikawa T. Ground-space bilateral teleoperation of ETS-VII robot arm by direct bilateral coupling under 7-s time delay condition. IEEE Transactions on Robotics and Automation. 2004;20(3):499–511.

11. Postolache O, Hemanth DJ, Alexandre R, Gupta D, Geman O, Khanna A. Remote monitoring of physical rehabilitation of stroke patients using IoT and virtual reality. IEEE Journal on Selected Areas in Communications. 2020;39(2):562–573.

12. Hojatmadani M, Rigsby B, Reed KB. Time Delay Affects Thermal Discrimination. IEEE Transactions on Haptics. 2022;15(2):451–457.

13. Ivanova E, Eden J, Zhu S, Carboni G, Yurkewich A, Burdet E. Short time delay does not hinder haptic communication benefits. IEEE Transactions on Haptics. 2021;14(2):322–327.

14. Melendez-Calderon A, Bagutti L, Pedrono B, Burdet E. Hi5: A versatile dual-wrist device to study human-human interaction and bimanual control. In: IEEE/RSJ International Conference on Intelligent Robots and Systems (IROS); 2011. p. 2578–2583.

15. Ivanova E, Carboni G, Eden J, Krueger J, Burdet E. For motion assistance humans prefer to rely on a robot rather than on an unpredictable human. IEEE Open Journal of Engineering in Medicine and Biology. 2020;.

16. Franklin DW, Burdet E, Tee KP, Osu R, Chew CM, Milner TE, et al. CNS learns stable, accurate, and efficient movements using a simple algorithm. Journal of Neuroscience. 2008;28(44):11165–11173.

17. Hasson CJ, Gelina O, Woo G. Neural control adaptation to motor noise manipulation. Frontiers in human neuroscience. 2016;10:59.

18. Börner H, Carboni G, Cheng X, Takagi A, Hirche S, Endo S, et al. Physically interacting humans regulate muscle coactivation to improve visuo-haptic perception. Journal of Neurophysiology. 2023;129(2):494–499.

19. Balasubramanian S, Melendez-Calderon A, Roby-Brami A, Burdet E. On the analysis of movement smoothness. Journal of Neuroengineering and Rehabilitation. 2015;12(1):112.

20. Kasuga S, Crevecoeur F, Cross KP, Balalaie P, Scott SH. Integration of proprioceptive and visual feedback during online control of reaching. Journal of Neurophysiology. 2022;127(2):354–372.

21. Oldfield RC. The assessment and analysis of handedness: the Edinburgh Inventory. Neuropsychologia. 1971;9(1):97–113.

22. Friston S, Karlström P, Steed A. The effects of low latency on pointing and steering tasks. IEEE Transactions on Visualization and Computer Graphics. 2015;22(5):1605–1615.

23. Pavlovych A, Stuerzlinger W. Target following performance in the presence of latency, jitter, and signal dropouts. In: Graphics Interface. vol. 2011; 2011. p. 33–40.

